# Investigating the Interactomic Landscape of Survival Motor Neurons (SMN) and the SMN Δ7 truncated protein

**DOI:** 10.1101/2025.08.11.669646

**Authors:** Bobby Beaumont, Silvia A. Synowsky, Sally L. Shirran, Judith E. Sleeman

## Abstract

This protocol describes a methodology combining TurboID - a recently developed proximity biotinylation technique - with conventional epitope-tag based co-immunoprecipitation (Co-IP) to analyse protein-protein interactions (PPIs) in cell culture systems. This integrated approach allows for the targeted examination of both transient and stable protein interactors, enhancing our understanding of protein dynamics.

TurboID captures transient interactions often missed by Co-IP, which captures high affinity, stable interactors. Combination of both techniques enables direct comparison of interaction strengths, providing insights into the dynamic nature of protein interactions within cells. The rapid biotinylation capability of TurboID reduces background noise and false positives while Co-IP enriches stable interactors, together improving data quality and interpretation of the interactomic landscape.

Demonstrating the efficacy of this methodology, proteins relevant to the pathology of Spinal Muscular Atrophy were utilised to explore variations at the interactome level. The use of identical starting cell lysates for both TurboID and Co-IP minimised variability and ensured datasets were comparable, allowing for consistency and enhancing the reliability of findings regarding the nature and strength of protein interactions. This novel framework effectively combines both innovative and classical techniques while maintaining consistency in sample handling, advancing our understanding of the intricate networks that govern cellular processes.

## Introduction

Interactomics is a branch of proteomics focussing on the study of interactions between molecules within an organism; it aims to characterize the complicated networks of protein-protein interactions (PPIs) that regulate cellular processes in both health and disease. PPIs are a crucial component of all processes in the cell - ranging from signal transduction to metabolic pathways - and they play a pivotal role in the organization and function of the proteome (Liu et al., 2024). These PPIs can be strong - involving stable complexes with specific binding sites - or transient, weak interactions, necessary for dynamic cellular processes (Mezentsev et al., 2022). Strong interactions are characterized by high affinity, specificity, and stability; weak interactions are more transient and dynamic, often influencing protein complex assembly and regulatory mechanisms during signal transduction (Weimer et al., 2023). Understanding the characteristics of both strong and weak PPIs is essential for unravelling the complexity of biological systems and the pathomechanisms of disease (Liu et al., 2024).

### Co-Immunoprecipitation

Various techniques to study PPIs exist, and one popular method is Protein Co-Immunoprecipitation (Co-IP). Co-IP is a classical technique in molecular biology and is used to identify protein interactors via pulldown of a target protein along with its binding partners from a mixed sample using antibodies (Lo Sardo, 2023). This method relies on the principle that by precipitating one protein, the other proteins which are in complex with it can also be precipitated. Co-IP has historically been important for studying *in vivo* protein-protein and protein-nucleic acid interactions, aiding in the identification of interacting molecules when combined with other methods including MS and Western Blot (reviewed in (Free et al., 2009)). Overall, Co-IP serves as a powerful tool for investigating strong protein interactions and aids in the understanding of complex protein networks.

### Proximity Biotinylation

Unlike classical Co-IP, TurboID is a more recent technique which labels the interacting partners of a protein of interest (POI) – also known as the bait protein - in a proximity-dependant manner (Figure 1). The TurboID protein is a modified BirA biotin ligase which, when conjugated to a protein, will biotinylate all proteins present within a 10-nanometer sphere upon the addition of biotin to culture media (Li et al., 2019). TurboID is a variation on the BioID (BirA) labelling technique, produced via targeted mutagenesis of the BirA ligase (Branon et al., 2018). This results in a variation of BirA which works more rapidly, reducing the time cells are exposed to biotin and thus resulting in fewer false positives and background noise that can occur in longer incubation times *in vitro*. Unlike Co-IP which reveals only stable and high affinity PPIs, TurboID can capture transient, weaker protein interactions (Samavarchi-Tehrani et al., 2020).

**Figure 1:**
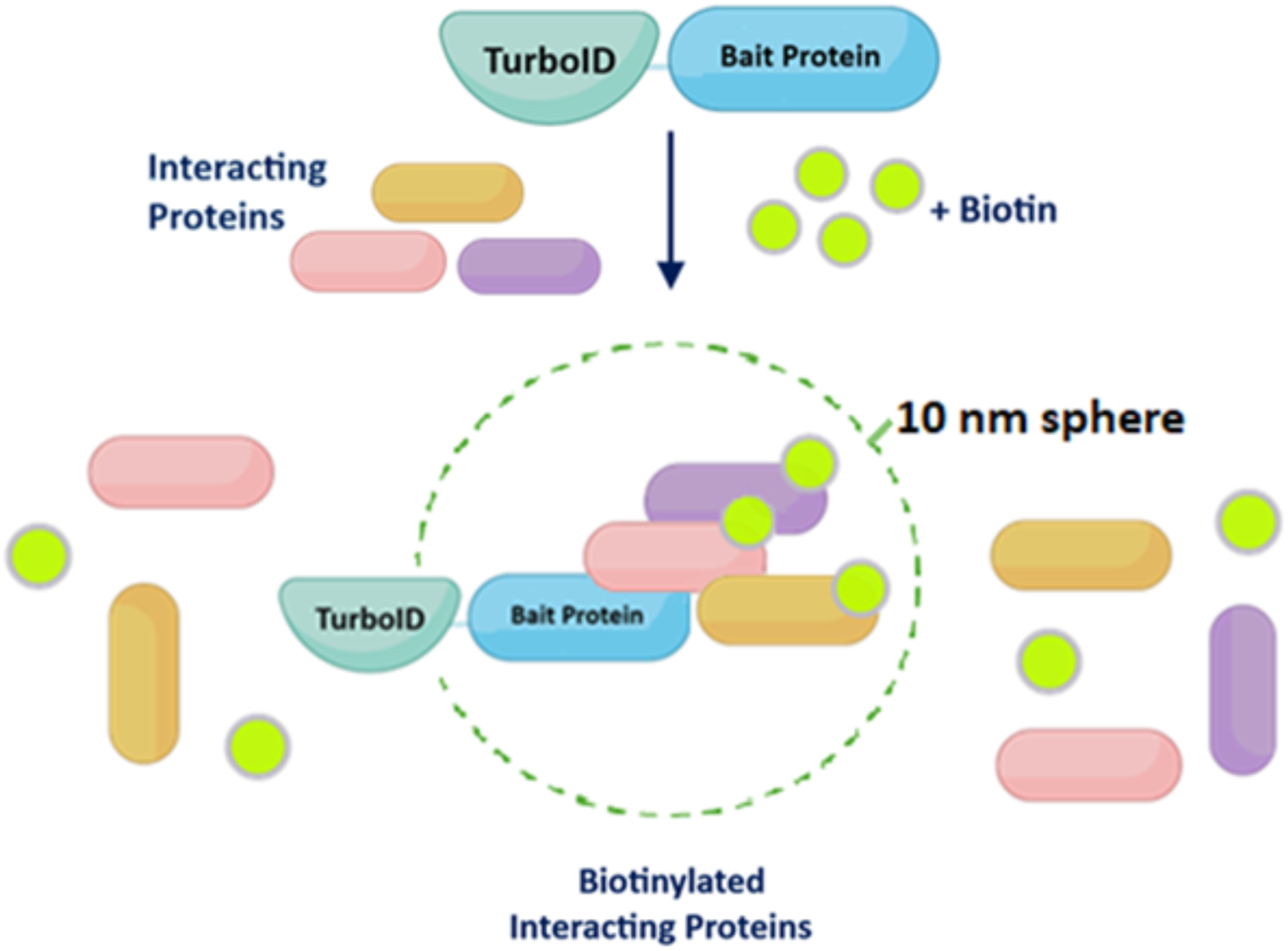
TurboID Captures Protein Interactors of Attached Bait Proteins/POIs. Upon the addition of biotin, interacting protein partners of a selected bait protein/POI fused to TurboID become biotinylated, allowing for efficient pulldown of tagged proteins using Streptavidin.

### Combinatorial Co-Immunoprecipitation and Proximity Biotinylation

As analysis of datasets derived from Co-IP and Proximity Biotinylation experiments can provide overlapping yet discrete information, the similarities and differences in these proteomes can be harnessed to further elucidate the nature of PPIs for a given bait protein. Combinatorial Co-IP and TurboID can be performed by adding an affinity tag to the TurboID-POI construct. While there are several affinity tags available (for an overview, see (**Kimple et al., 2013**)), for the purposes of this methodology, the FLAG tag was chosen to represent the Co-IP/Affinity Purification technique.

The FLAG tag is a short peptide sequence consisting of eight amino acids (DYKDDDDK) that can be fused to proteins for specific downstream applications in molecular biology. Adding a FLAG tag to a protein enables specific detection and study of the tagged protein of interest via antibody-based detection methods, using antibodies against the epitope tag (Hopp et al., 1988). FLAG-tagged proteins - along with their interacting protein partners - can thus be separated from complex protein mixtures and analysed by mass spectrometry (Fedorova & Dorogova, 2019). By tagging the TurboID-POI construct with a FLAG tag, concurrent Co-IP can be performed on TurboID-transduced stable cell lines to give complementary proteomic datasets.

Utilising these techniques in tandem makes it possible to design an experimental workflow that can produce datasets elucidating which interactions are consistent (or differ) between both methodologies (Figure 2). The Co-IP method produces data enriched in stable, high affinity protein interactors which can be compared to the data produced by TurboID-induced biotinylation; this data will include not only the stable interactors, but also lower affinity, transient proteins of interest. Simultaneous culture of cells in SILAC (Stable Isotope Labelling by Amino Acids in Cell Culture) media allows for the production of ratios between different SILAC conditions (the Light/Medium/Heavy labels) thus assigning quantitative values to mass spectrometry data (Chen et al., 2015). The method described uses the same starting cell lysates and the same wash conditions for both techniques, minimising the passage-to-passage variation that can occur during cell culture and the variability observed in differing wash stringencies.

**Figure 2:**
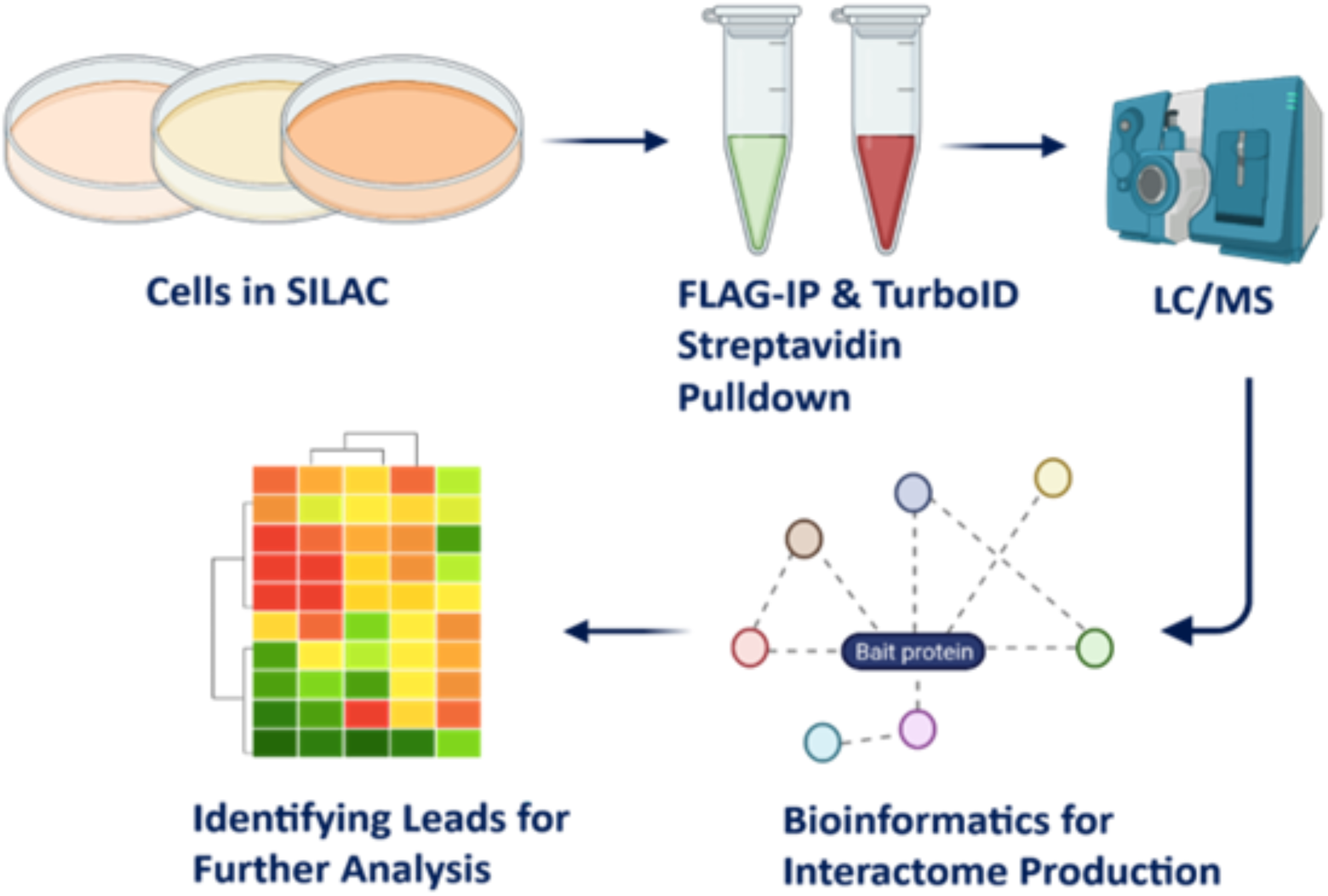
Experimental workflow to identify interacting partners of specific Bait Proteins using Complementary Co-IP, TurboID and Mass Spectrometry. A graphical overview of the experimental workflow, utilising stable cell lines expressing FLAG-TurboID-POI for FLAG-Immunoprecipitation and Streptavidin Pulldown. Liquid Chromatography/Mass spectrometry (LC/MS) of SILAC-labelled samples and subsequent data analysis of raw MS files allows for production of quantitative interactomes and identification of novel protein interactors.

## Materials and Reagents

- Cell Lines and Culture Media

- Cell lines of interest
- DMEM (Corning) [Cat No: 10-013-CV]
- Foetal Bovine Serum (FBS, Gibco) [Cat No: 10270-106]
- Penicillin/Streptomycin (Thermo) [Cat No: 15140122]
- Phosphate-Buffered Saline (PBS)
- Transfection reagent (eg Effectene or Transduction reagent (eg PEI)
- TurboID plasmids
- Fixatives and Staining Reagents

- Paraformaldehyde (PFA, Sigma) [Cat No: P6148]
- Triton-X100 (Sigma) [Cat No: X100]
- Goat Serum (Thermo) [Cat No: 16210064]
- DAPI (Sigma) [Cat No: D9542]
- Prolong Gold Antifade Reagent (Life Technologies) [Cat No: P36930]
- Doxycycline (VWR) [Cat No: 101515-454]
- Biotin (ThermoFisher) [Cat No: 29129]
- 488-Streptavidin (Licor) [Cat No: 926-32230]
- Primary Antibodies
- Protein Analysis Reagents

- SDS-PAGE gel components (Acrylamide, dH2O, Rx Buffer pH 8.8, Rx Buffer pH 6.8, TEMED, 10% APS)

- Acrylamide (Sigma) [Cat No: A3553]
- TEMED (Sigma) [Cat No: T9281]
- 10% APS (Sigma) [Cat No: A3678]
- Amersham Protran Nitrocellulose membrane (GE Healthcare) [Cat No: 10600002]
- Ponceau solution (Sigma) [Cat No: P7170]
- Milk Powder
- PBS-T (1% Tween-20 in PBS)
- NuPAGE™ LDS Sample Buffer (4X) (Thermofisher) [Cat No: NP0007] PageRuler™ Plus Prestained Protein Ladder, (Thermofisher) [Cat No: 26619]
- SILAC Media Components

- DMEM for SILAC (Thermo Scientific) [Cat No: 88364]
- Dialyzed FBS (Gibco) [Cat No: A3382001]
- Leucine (Sigma) [Cat No:[L8912]
- R0 Arginine (Sigma) [Cat No: A5006]
- K0 Lysine (Sigma) [Cat No: L8662]
- R6 Arginine (Sigma) [Cat No: 60803]
- K4 Lysine (Sigma) [Cat No: 60841]
- R10 Arginine (Sigma) [Cat No: 60801]
- K8 Lysine (Sigma) [Cat No: 60842]
- Mass Spectrometry and Sample Preparation Reagents

- Low Salt RIPA buffer (50 mM Tris, pH 7.5, 150 mM NaCl, 1% NP-40, 0.5% deoxycholate, protease inhibitor tablet)
- Enzyme-Free Cell Dissociation Buffer (Gibco) [Cat No: 13151014]
- Sepharose beads (ThermoFisher) [Cat No: 101970]
- Protein A Beads (Thermo) [Cat No: 20333]
- Invitrogen Mouse Anti-FLAG antibody (Invitrogen) [Cat No: MA1-91878]
- 4X (LDS) Sample Buffer (NuPage) [Cat No: NP0007]
- DTT (VWR) [Cat No: 97061-056]
- Ammonium bicarbonate (Thermo) [Cat No: 09830]
- Magnetic Streptavidin-Agarose Beads (Pierce) [Cat No: 88817]
- AF800-Streptavidin (JIR) [Cat No: 016-440-084]
- Trypsin (Sigma) [Cat No: T4799]
- LC-MS reagents (Pepmap100 C18, Trifluoroacetic Acid (TFA), acetonitrile, formic acid)
- MaxQuant software (https://www.maxquant.org/)
- Perseus Software (https://maxquant.net/perseus/)
- Microsoft Excel

### Equipment

- Magnetic rack
- Centrifuge
- End over end rotator
- Speedvac
- Western Blot Imaging System eg Licor Odyssey CLx
- Sonicator
- Biophotometer
- Semi-Dry Blotter

## Method

### Plasmid Design and Generation

1. Plasmids were generated according to design instructions supplied to Vector Builder. However, TurboID plasmids can be purchased from AddGene (FLAG-TurboID is Addgene Plasmid #124646)
2. The addition of alternative affinity tags (and other features) can be performed via in-house cloning or via an external agency (e.g. Vector Builder)
3. For best results, the affinity tag should be placed at the TurboID N-terminus (resulting in plasmids with the following structure: Tag-TurboID-POI. A 5xFLAG tag was utilised in the experiments described below).
4. A separate control plasmid consisting of Tag-TurboID but no bait protein is necessary to compare background levels of biotinylation.
5. Utilisation of a Tet-On system ensures that the TurboID construct is expressed only in the presence of Doxycycline; this reduces potential off-target effects or toxicity which may be present in constitutive expression) (May et al., 2020). In the data discussed below, a two-step Tet-On system was utilised; this included transduction of a Tet-On Helper plasmid containing silencer elements into a parental cell line, which – post-selection – were then transduced with the Flag-TurboID Plasmids.
6. An appropriate antibiotic resistance cassette is required for selection of transduced cells: a puromycin selection cassette may be utilised for fast (5-10 days) selection post-transduction/transfection.
7. Consider the addition of a fluorescent live cell marker protein such as mKate2 to allow for easier determination of transduction/transfection efficiency (Shcherbo et al., 2009).
8. While the above features can be designed and encoded in traditional mammalian plasmids, the following protocol focuses on the use of Lentiviral plasmids.

### Generation of Human Stable Cell Lines

1. Culture cells in optimal media of choice for the specific cell line (e.g. for Hela cells, DMEM supplemented with 10% FBS and 1% penicillin/streptomycin).
2. Incubate cells at 37°C and 5% CO2.
3. Passage cells routinely upon reaching ∼70% confluency at a 1:10 ratio to maintain health and viability.

### Cell Transduction and Selection

1. Seed 8.5 x 10⁵ HeLa cells into T25 flasks.
2. Incubate the cells for 24 hours.
3. After 24 hours, replace the media with the serum-free equivalent.
4. Transduce the cells with a volume of either FLAG-TurboID, FLAG-TurboID-SMN, or FLAG-TurboID-SMNΔ7 viral supernatant, equivalent to an MOI of 10.
5. After 6 hours of transduction, add 20% FBS-supplemented media to the flask.
6. Incubate the cells for 72 hours post-transfection.
7. Transfer the cells to new T25 flasks for selection using puromycin at a concentration of 200 ng/ml. Note: determine the optimal antibiotic level required per cell line by calculating a kill curve.
8. Replace the selection medium every 2 to 3 days for a duration of 5 to 10 days.
9. After selection, transfer the cells to T75 flasks for expansion. TurboID Induction of Protein Expression
10. After expansion of cell lines post-selection, seed transduced cells onto coverslips in a multiwell plate and induce with 1-2 µg/ml doxycycline.
11. Incubate cells for 48 hours at 37°C.
12. If cells contain a fluorescent marker, transduction efficiency can be calculated via imaging on a fluorescent microscope prior to biotin incubation.
13. Incubate cells with 150 µM biotin for 1 hour at 37°C. *Note*: empirically determine the optimal biotin concentration and timepoint for each transduced cell line. A working range of 50 – 300 µM biotin concentration and several timepoints to a maximum of 1 hour will provide a solid overview.
14. Fix cells and stain with 488-Streptavidin and POI-specific primary antibodies as described below.

### Cell Staining

1. Wash coverslips in PBS.
2. Add 5 ml of 3.7% paraformaldehyde fixative buffer to a 10 cm dish containing coverslips and incubate for 10 minutes at room temperature.
3. Remove PFA and wash coverslips twice with PBS, then store in PBS at 4°C until use.
4. Permeabilize cells with PBS/0.1% Triton-X100 on a shaker for 15 minutes.
5. Block coverslips with PBS/1% goat serum for 20 minutes.
6. Incubate coverslips in primary antibodies against the POI, diluted with PBS and 1% Goat Serum for 1 hour in a humidity chamber.
7. Wash coverslips three times in PBS, 10 minutes each time.
8. Incubate coverslips in secondary antibodies (including fluorescent streptavidin) in PBS and 1% Goat Serum for 1 hour. Note: ensure that the secondaries do not overlap with any fluorescent marker proteins.
9. Wash once in PBS for 10 minutes.
10. Counter-stain coverslips with DAPI (1.6 µg/ml) for 10 minutes.
11. Wash coverslips in dH2O for 10 minutes.
12. Mount coverslips onto slides using 8 µl of Prolong Gold Antifade Reagent. Allow 24 hours for curing in the dark.
13. Image cells in the appropriate channels using a fluorescent microscope. Analyse the localisation of the TurboID bait protein to ensure it localises to the same cellular locations as the native protein.

### SDS-PAGE

1. To harvest cells, dissociate cells using Trypsin/EDTA and spin at 200 RCF for 5 minutes; discard media and resuspend in ice-cold PBS. Repeat this for a total of 3 washes.
2. Resuspend cell pellets in up to 1 ml of fresh Lysis Buffer.
3. Sonicate on ice at 5 microns for 3 x 15 seconds, with 10 seconds of rest on ice between.
4. Measure total protein via Bradford Assay using a Biophotometer. Flash freeze lysates in Liquid N^2^ and store at −80°C until ready for use.
5. Prepare 10% SDS-PAGE gels or use NuPAGE 4-12% pre-cast gels.
6. Add 5 ul of 4X loading dye to 15 ul of lysate and heat to 95°C for 10 minutes.
7. Load 40 µg of protein alongside a molecular weight marker ladder.
8. Run hand-poured gels for 80 minutes at 150V or NuPage pre-cast gels for 45 minutes at 200V.

### Western Blotting

1. Transfer proteins to a nitrocellulose membrane using a Turbo-Blot semi-dry blotter.
2. Post-transfer, stain membrane with ponceau solution for 5 minutes, wash with dH2O, scan, and de-stain with PBS.
3. Block membrane in 5% milk powder and PBS for 1 hour at 4°C.
4. Incubate membrane in primary antibodies overnight at 4°C in 5% milk and PBS-T.
5. Wash membrane three times for 5 minutes in PBS-T.
6. Incubate membrane in secondary antibodies in 5% milk powder and PBS-T for 1 hour at room temperature.
7. Wash membrane three times for 5 minutes each in PBS-T and image using the LiCor Odyssey® CLx imaging system.
8. Examine expression of the POI using specific primary antibodies. Similarly, examine levels of biotinylation utilising a fluorescent streptavidin secondary antibody. Check for specific expression of the TurboID construct using primary antibodies against the affinity Tag (e.g. anti-FLAG antibody).

### SILAC Cell Culture

1. Prepare Light, Medium, and Heavy SILAC media as described in Table 1.
2. Culture cells for more than 3 passages in corresponding media. To produce enough protein, it is recommended to harvest at least 3 x T150 flasks per cell line.
3. Obtain a minimum of 2 biological replicates per cell line, switching labels between conditions (e.g if Cell line X in Bio Rep 1 is grown in Light SILAC media, in Bio Rep 2, Cell line X will be cultured in Medium SILAC Media)
4. To harvest, dissociate cells using Enzyme-Free Cell Dissociation Buffer, spin at 200 RCF, discared media and resuspend in ice-cold PBS. Repeat this for a total of 3 washes.
5. Flash freeze cell pellets in liquid nitrogen before further processing.

### SILAC Sample Preparation for Mass Spectrometry Analysis Cell Lysis

1. Resuspend cell pellets in up to 1 ml of fresh Low Salt RIPA buffer.
2. Sonicate on ice at 5 microns for 3 x 15 seconds, with 10 seconds of rest on ice between.
3. Measure total protein via Bradford Assay using a Biophotometer.
4. Store lysates at −80°C.

### FLAG Immunoprecipitation (Co-IP)

1. Pre-clear 2 mg of input protein in up to 0.7 ml of Low Salt Ripa Buffer with 20 µl of Sepharose beads via gentle rotation (15 RPM) for 30 min at 4°C using an end-over-end rotator.
2. Pellet beads and save 40 µg of the supernatant as the “Precleared” sample.
3. Add remaining supernatant to either 30 µl Protein A Beads and 2 µg/ml Invitrogen Mouse Anti-FLAG antibody (Antibody sample) or 30 µl Protein A Beads only (No Antibody sample).
4. Incubate samples at 4°C overnight with gentle rotation (15 RPM).
5. Pellet beads and save 40 µl of the supernatant as the “Bead Unbound” sample.
6. Wash beads in 500 µl of Low Salt RIPA buffer four times.
7. For western blotting, add 4X LDS Sample Buffer with DTT to a final concentration of 50 mM and heat beads for 25 mins at 65°C.
8. For on-bead digestion and mass spectrometry, wash combined bound bead fractions (Heavy/Medium/Light) five times with 50 mM ammonium bicarbonate solution.

### Streptavidin Pulldown

1. Pre-clear 2 mg of input protein in up to 0.7 ml of Low Salt Ripa Buffer with 20 µl of Sepharose beads via gentle rotation (15 RPM) for 30 min at 4°C using an end-over-end rotator.
2. Pellet beads and save 40 µl of the supernatant as the “Precleared” sample.
3. Wash Magnetic Streptavidin-Agarose Beads three times in Low Salt RIPA buffer.
4. Add 30 µl of beads to remaining precleared cell lysates and mix overnight at 4°C with gentle rotation.
5. Pellet beads using a magnetic rack and save 40 µl as an “Unbound” sample.
6. For western blotting, wash beads four times with Low Salt RIPA buffer and elute proteins using 30 µl of elution buffer (30 mM Biotin in 2% SDS) and 20 µl of 4X LDS Buffer with 50 mM DTT.
7. Heat beads for 25 mins at 65°C to reduce proteins.
8. For on-bead digestion and mass spectrometry, combine the bound bead samples (Heavy/Medium/Light) and wash five times with 50 mM ammonium bicarbonate solution.

### Mass Spectrometry Analysis

**Note: Specific MS methodology will depend on the instrumentation available. The following is optimised for the Orbitrap Fusion Lumos.**

1. Perform trypsin digestion of beads overnight at 37°C.
2. Reduce final sample volume to 20 µl using a Speedvac.
3. Subject peptides to LC-MS/MS using an Ultimate 3000 RSLC coupled to an Orbitrap Fusion Lumos mass spectrometer with a FAIMS interface.
4. Inject peptides onto a reverse-phase trap for pre-concentration and desalting.
5. Elute peptides from the analytical column using a linear solvent gradient.
6. Operate mass spectrometer in Data Dependent Acquisition (DDA) positive ion mode with a cycle time of 1.5 seconds.
7. Select peptides with charge states 2 to 5 for fragmentation using HCD as collision energy.

**Table 1:** SILAC Media Recipes. Components required for the creation of 500mls of SILAC Light (R0K0) (yellow cells in table), Medium (R6K4) (blue cells in table) and Heavy (R10K8) (green cells in table) media.

| Condition | Media | Components |
| --- | --- | --- |
| **Light** | DMEM for SILAC | - 10% dialyzed FBS |
| (R0K0) |  | - 1% Penicillin/Streptomycin (P/S) |
|  |  | - 1mL Leucine (52mg/ml) |
|  |  | - 0.5mL R0 arginine (84mg/ml) |
|  |  | - 0.5mL K0 lysine (146mg/ml) |
| **Medium** | DMEM for SILAC | - 10% dialyzed FBS |
| (R6K4) |  | - 1% Penicillin/Streptomycin (P/S) |
|  |  | - 1mL Leucine (52mg/ml) |
|  |  | - 0.5mL R6 arginine (84mg/ml) |
|  |  | - 0.5mL K4 lysine (146mg/ml) |
| **Heavy** | DMEM for SILAC | - 10% dialyzed FBS |
| (R10K8) |  | - 1% Penicillin/Streptomycin (P/S) |
|  |  | - 1mL Leucine (52mg/ml) |
|  |  | - 0.5mL R10 arginine (84mg/ml) |
|  |  | - 0.5mL K8 lysine (146mg/ml) |

### Data Processing

1. Search raw MS data files against the UniProt Human Proteome database using MaxQuant software (https://www.maxquant.org/)
2. Set peptide tolerance to 10 ppm and select trypsin as the enzyme.
3. Select fixed modification as carboxyamidomethylation of cysteine.
4. Select variable modifications as oxidation of methionine and N-terminal acetylation.
5. Select SILAC labels of Arg6 (R6), Arg10 (R10), Lys4 (K4), and Lys8 (K8).
6. Choose a minimum of 1 unique peptide, minimum ratio count of 2, and quantitation based on razor and unique peptides.
7. Set peptide and protein FDR to 0.01.
8. Run the analysis.

### Data Analysis

- Process output data using the ProteinGroups file in Microsoft Excel or Perseus to produce ratios, proportions, and percentages.
- Account for relevant controls and remove Control-only protein hits from datasets.
- Charts can be produced using GraphPad Prism 6 or Microsoft Excel.
- Perform statistical analysis in either Microsoft Excel or Perseus software (https://maxquant.net/perseus/)
- Select proteins with significant changes in interactivity (P Value < 0.05 and Fold Change
- < 0.5 and > 1.5) for further analysis.
- Compare proteomes of Co-IP vs Streptavidin Pulldown to label protein interactors as either stable (present in both Co-IP and Streptavidin Pulldown) or potentially transient (present in Streptavidin Pulldown only).

## Results and Discussion

The above methodology was used to study the differential interactomes of proteins relevant to the pathology of Spinal Muscular Atrophy (SMA). SMA is a genetic neuromuscular disorder caused by mutations in the SMN1 gene, resulting in the lack of production of the SMN protein (Lefebvre et al., 1995). Humans possess a backup SMN gene, known as SMN2, which – due to a splicing error – produces insufficient amounts of full-length SMN protein. The primary gene product of SMN2 is SMNΔ7, an SMN isoform which lacks the Exon 7 of full-length SMN and is generally considered non-functional and unstable (Lefebvre et al., 1997).

Utilising lentiviral constructs encoding FLAG-TurboID-SMN or FLAG-TurboID-SMNΔ7, stable Hela cell lines were generated, harvested and processed as described to probe the differential interactomes between the two protein isoforms (for plasmid maps, see Supplementary Information). Localisation of each protein construct was investigated via immunostaining and examination of biotinylation patterns were performed simultaneously (Figure 3A). Each of the Hela TurboID cell lines demonstrated sufficient levels of biotinylation in all three experimental conditions (TurboID-Control, TurboID-SMN and TurboID-SMNΔ7) when induced for 48 hours with 1ug/ml of doxycycline.

**Figure 3:**
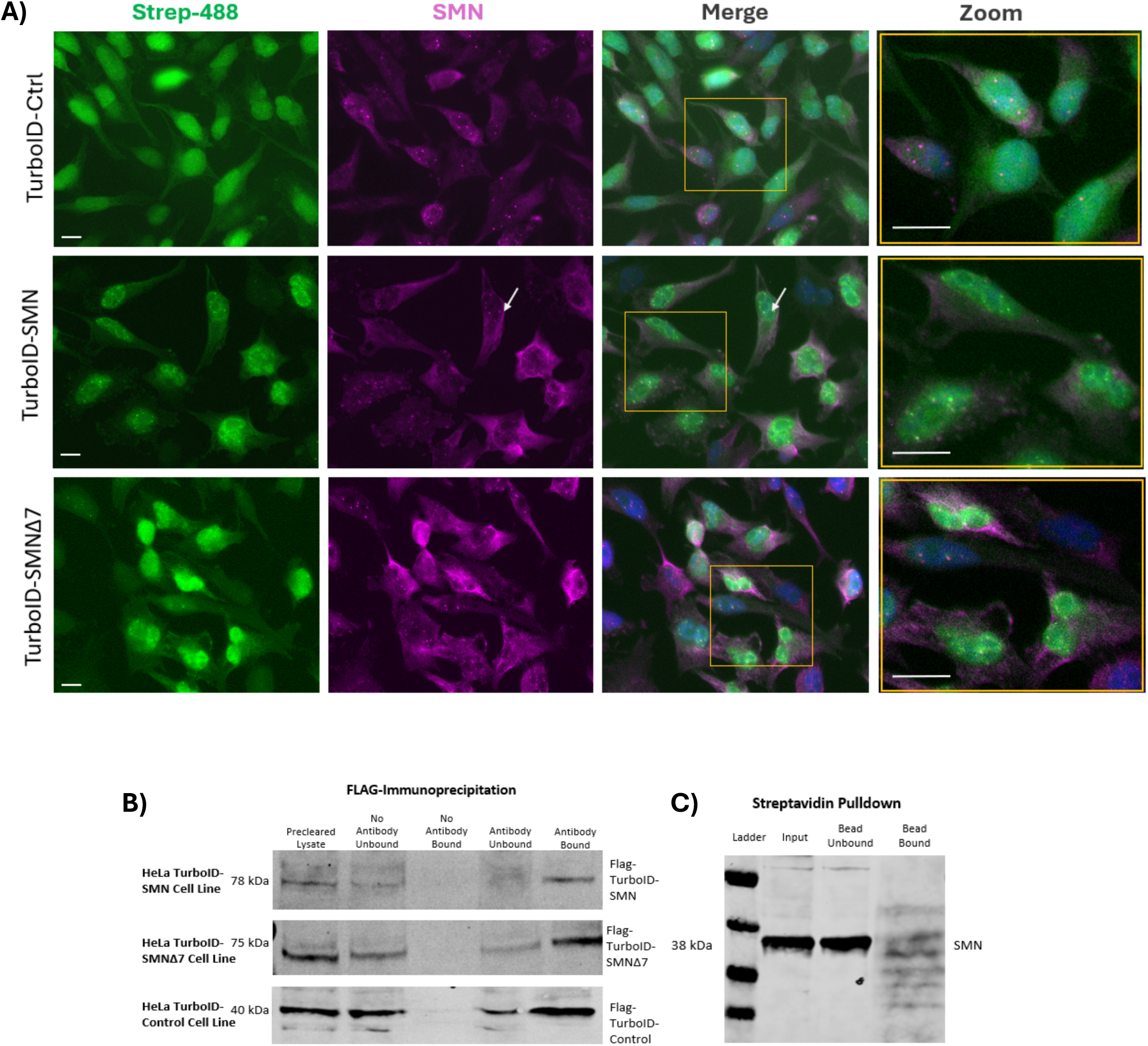
Validation of Hela Turbo-ID stable cell lines confirms Bait Proteins localise correctly, biotinylate interacting proteins and pull down efficiently. **A)** Representative fluorescent imaging of 48hr induced (1ug/ml doxycycline) and 1 hr biotin incubation (150uM) TurboID Hela cell lines. TurboID-Control (top), TurboID-SMN (centre) and TurboID-SMNΔ7 (bottom) cells were stained with Streptavidin-488 (Licor) (Green) to indicate biotinylation and mouse Anti-SMN (BD Biosciences) (Magenta). Nuclei were counterstained with DAPI (blue). Scale = 10 microns. Images were obtained on the EVOS M5000 Imaging System at a magnification of 40x. White arrows indicate nuclear SMN-containing gems of Cajal Bodies. **B)** FLAG Immunoprecipitation of Hela TurboID cell lines. Immunoprecipitation was performed as described with Protein A beads and a mouse Anti-FLAG antibody (Invitrogen). The “No Antibody” lanes represent the negative controls i.e. lysates were incubated with beads only. The “Antibody” lanes indicate that lysates were incubated with both beads and antibody. Starting input = 2mg of cell lysate. Blots were probed with a rabbit Anti-FLAG (Sigma) antibody. **C)** Streptavidin Pulldown of a TurboID cell lysate using Magnetic Streptavidin-Agarose Beads exhibited biotinylation on Western Blot after staining with Streptavidin-680. Lysate derived from 48hr dox-induced (1ug/ml) TurboID-Control Hela cell line incubated with 150uM of Biotin for 1hr at 37 degrees Celsius prior to harvest. Blots were probed with Streptavidin-680 (JIR) and mouse Anti-SMN (BD Sciences). Starting input = 2mg of cell lysate.

Biotinylation with 150uM of biotin for 1 hour at 37 degrees Celsius gave a strong, specific biotinylation signal consistent the with signal expected for each cell line’s transduced construct. The biotinylation signal observed in the TurboID-Control cell line was consistent with a diffuse, non-specific cellular localisation. The TurboID-SMN line demonstrated biotinylation consistent with nuclear SMN gem staining (see white arrows in Figure 3A, middle row) as well as a level of diffuse, cytoplasmic staining whereas the biotinylation present in the TurboID-SMNΔ7 line was primarily concentrated in the nucleus, consistent with expected SMNΔ7 expression.

FLAG Immunoprecipitation and Streptavidin Pulldown capabilities were examined by western blot prior to final cell culture in SILAC (Fig 3B and C). Immunoprecipitation reactions using 2ug/ml of Invitrogen Mouse Anti-FLAG antibody were successfully performed and resulted in protein bands at the expected locations in western blot.

Utilisation of a low salt (150mM) RIPA buffer as both cell lysis buffer and wash buffer, and elution conditions optimal for the Protein A beads resulted in efficient capture of the TurboID proteins. The same lysis and wash conditions were utilised for the Streptavidin Pulldowns, which were instead performed with Magnetic Streptavidin Beads. In addition to being probed for SMN, these immunoblots were also stained with fluorescent streptavidin to confirm efficient biotinylation. Once blots and microscopy confirmed correct expression and localisation, respectively, of the TurboID cell lines, cells were cultured in SILAC media prior to MS, to allow for quantitative comparison between interactomes.

The Co-IP/FLAG-IP method yielded fewer protein interactors (480 proteins) in comparison to the number identified by Streptavidin Pulldown (2264 proteins) by MS. This almost 5 times larger interactome in the Streptavidin Pulldown was to be expected, given that it includes transient, weak interactors in addition to strong interactors. Of the proteins identified via Co-IP, 75% of these interactors were also present in the Streptavidin Pulldown, confirming that these were indeed stable interacting partners of SMN or SMNΔ7.

Both datasets were probed for the presence of common SMN interactors as derived from high confidence experimental data obtained from String-DB (Figure 4). The FLAG IP dataset contained 5 out of 12 common SMN interactors (42%) whereas the Streptavidin Pulldown contained 11 out of 12 (92%) of these proteins. Notably, the only protein not identified in the Streptavidin Pulldown data was the Small Nuclear Ribonucleoprotein E (snRNP E) which was identified in the FLAG IP dataset.

**Figure 4:**
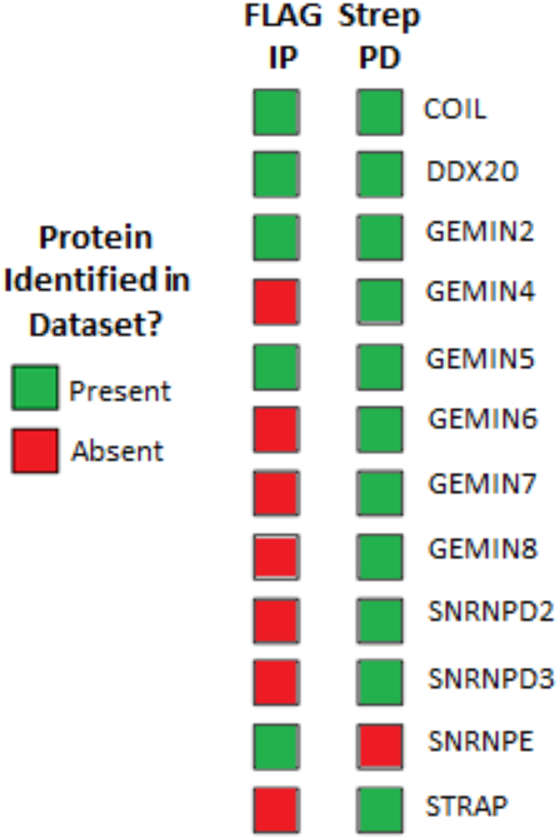
Both Flag IP and Streptavidin Pulldown Datasets Contain common SMN interacting proteins, however Streptavidin Pulldown has a Significantly larger coverage.

Twelve high confidence (> 0.7 confidence score), experimentally derived SMN interacting proteins were selected from String-DB and the produced Mass Spectrometry datasets for each interactomic method were probed for their presence. Gene Names for these protein interactors are found to the right of the figure, whilst the proteins’ presence in the either the FLAG IP dataset (left column) or Streptavidin Pulldown (right column) is shown as a red box if absent in the dataset or a green box if present. Figure key is found to the left.

Using these datasets, further statistical and bioinformatic analysis allowed for production of lists of differentially interacting proteins specific to both FLAG-TurboID-SMN and FLAG-TurboID-SMNΔ7. To more closely examine the interaction patterns of the proteins differentially interacting with FLAG-TurboID-SMNΔ7 and FLAG-TurboID-SMN, volcano plots were created for each technique (Figure 5).

**Figure 5:**
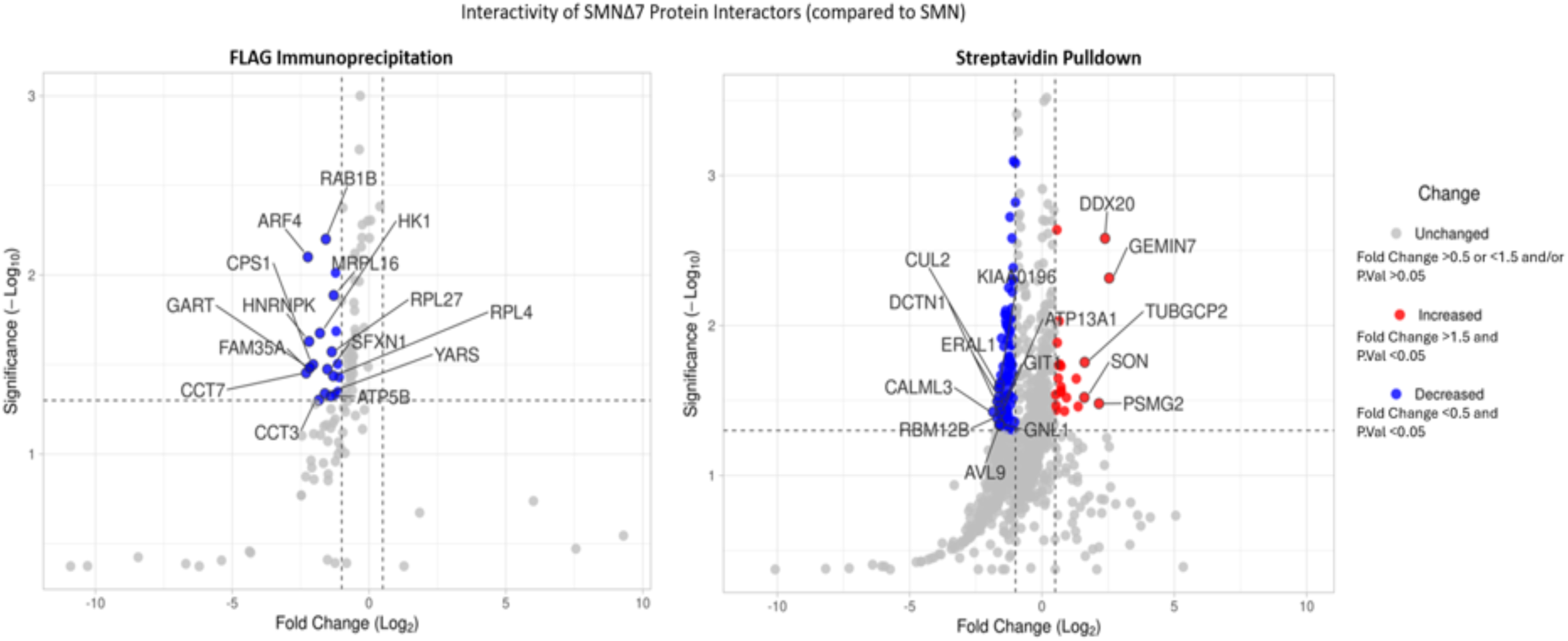
The Streptavidin Pulldown Data demonstrates Greater Interactome coverage than FLAG IP. Volcano plots showing differentially regulated proteins of interest in the Δ7 v SMN condition. Both the FLAG Immunoprecipitation data (Left) and the Streptavidin Pulldown data (Right) are displayed. Fold change (FC) cutoffs were set to show proteins with only FCs above 0.5 and below −1 (the log2 equivalent of an FC <0.5 and >1.5) The significance cutoff was set to 1.3 (the log10 equivalent of a 0.05 cutoff). The top 15 hits - ranked by the largest FC in either direction - were labelled with their gene name. Directionality of FC indicated by the colours defined in the key (found to the right of plots). Note: interactive versions of these plots are available as supplemental data figure S2.

Both plots display a similar profile insofar as the majority of significantly interacting proteins do not reach the cut off value (<0.5 or >1.5 Fold Change) necessary for highlighting. Interestingly, the FLAG IP data does not contain any proteins with increased interactivity which fit the parameters for inclusion in the graph; only decreased interactors attain the level of significance and meet the fold change cutoff. This is consistent with the literature describing SMNΔ7 as non-functional i.e. displaying reduced interactivity with canonical SMN interactors (Le et al., 2000). The Streptavidin Pulldown dataset, on the other hand, reveals a much wider scope of interactors – both increased and decreased in the FLAG-TurboID-SMNΔ7 condition compared to FLAG-TurboID-SMN. Of note, only ∼25% of the significantly differential (P<0.05) interactors in the FLAG IP condition were concordant with the significant interactors identified in the Streptavidin Pulldown. While there is a high level of overlap (75%) between identified proteins in both conditions, at the level of significance, there is evident variability between techniques. Whether this is primarily due to the greater number of identified interactors in the Streptavidin Pulldown remains an open question.

Given the mechanism of action of TurboID as a biotin ligase, however, it is important to note that there must be a measure of discretion when interpreting results; the biotinylation and enrichment of proteins via pulldown comprise interactors present in a 10-nanometre radius of the bait protein (Li et al., 2019). While the proteins may be in the right place at the right time, it does not necessitate that there is a direct interaction between the two. Hence it is important to apply biological context and previous knowledge when interpreting TurboID datasets.

In light of this, to confirm that the increased interactions observed in the Streptavidin Pulldown dataset were not due to the predominantly nuclear localisation of SMNΔ7, further bioinformatic analysis was performed to assign protein localisation data to each of the significantly increased and decreased interactors (Figure 6). From the increased interactor data, it can be concluded that these interactions are not solely due to the nuclear localisation of SMNΔ7; only one protein with increased interactivity is nucleus-specific (Protein SON) while six increased interactors are located only in the cytoplasm (DNAJC13, KIF5B, PFAS, TAF6, TUBGCP2 and UBAP1) and four localise to both compartments (GEMIN7, PRKACA, GEMIN4 and DDX20). Similarly, the localisation of several of those proteins with a *decreased* interaction with SMNΔ7 are nuclear-localised; this would not be the case if these were proteins TurboID-SMNΔ7 tagged purely based on proximity within the nucleus.

**Figure 6:**
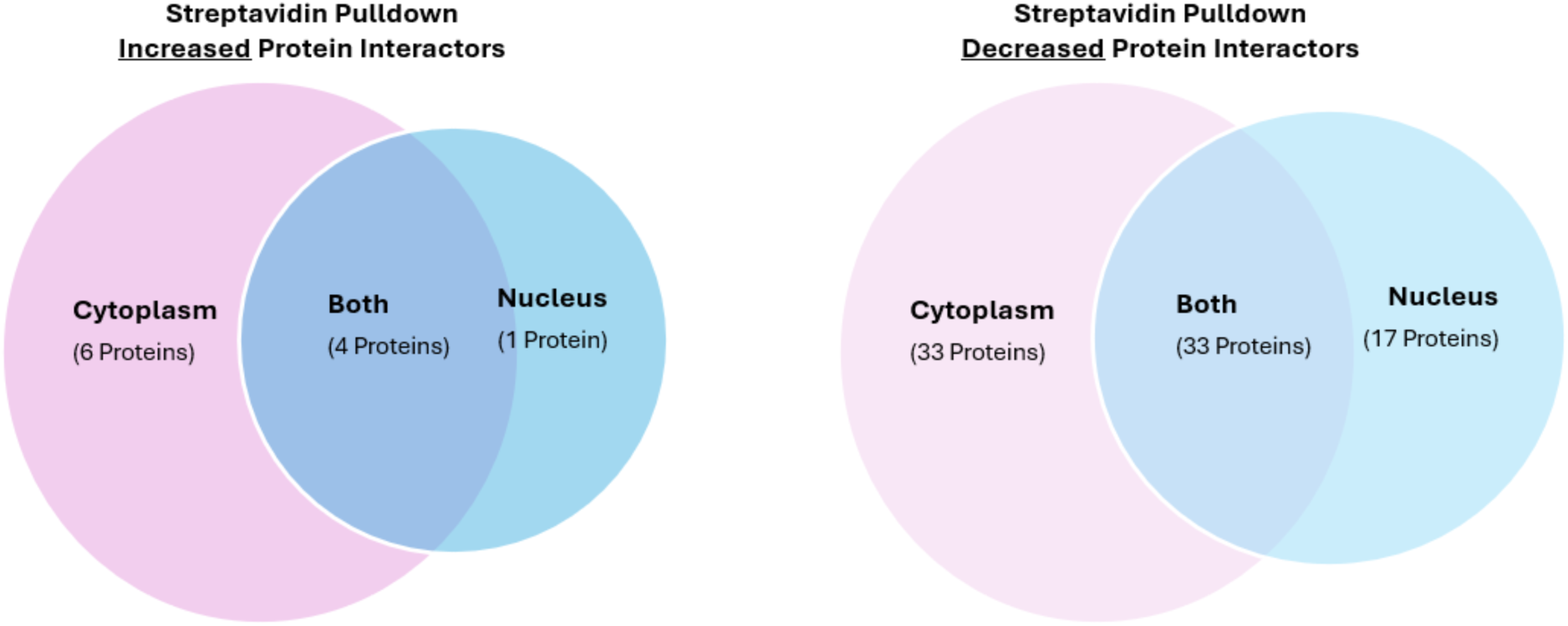
The Sublocalisation of Significant Protein Interactors Identified in the TurboID-SMNΔ7 Streptavidin Pulldown is not Confined to the Nucleus. Venn diagrams showing the cellular compartment (Cytoplasm/Nucleus/Both) of the proteins with increased interactions (left) and decreased interactions (right) in the Streptavidin Pulldown SMNΔ7 dataset, as compared to SMN (P<0.05; <0.05 FC >1.5)

While SMA is considered a neuronal disease, Hela cells provide an easy and efficient cell culture system to study protein-based interactomics. In previous SMA proteomics studies using human neuronal cells, negligible numbers of the differentially regulated proteins altered in SMA were specific to or enriched in neuronal cell lines. In a study using induced pluripotent stem cell (iPSC) motor neurons derived from patients with SMA type 1, none of the significantly altered proteins identified in SMA were either specific to neurons or enriched within neurons, when compared to a list of 130 neuronally-enriched proteins derived from the Human Protein Atlas (see Supplementary Data 2) (Fuller et al., 2015). Similarly, in a multi-timepoint study of human neuronal iPSCs derived from SMA patients, 3 of these neuronally enriched proteins were identified across numerous timepoints (Varderidou-Minasian et al., 2021). Only the neuron-enriched Ralyl protein was differentially expressed in SMA patients consistently, in 8 out of 10 timepoints. GNB1 was identified at two timepoints, whereas GAP43 was identified only once. Together, these data suggest that while SMA is a neuronal disease, a non-neuronal cell line (such as Hela) is an acceptable medium of study.

A loss of binding to canonical SMN interactors by SMNΔ7 is assumed to be the primary causative factor for its inability to recapitulate the function of SMN in SMA patients (Lefebvre et al., 1997).The data generated from the above comparative interactomics, however, suggests that many of the interactions in the SMNΔ7 isoform are not completely lost but simply decreased. There is a significant decrease observed in the interactivity between SMNΔ7 and several proteins (as illustrated most clearly in the Streptavidin Pulldown dataset), suggesting that it is not a complete loss of interactivity that hinders the functionality of SMNΔ7. Loss of functionality in SMNΔ7 may be due to a shift in the isoform’s axis of interactivity, with lower baseline interactions with several SMN specific interactors and, interestingly, increased interaction with others such as Gemins 4 and 7, two proteins which localise to both the nucleus and the cytoplasm (Cauchi, 2010). These - along with DDX20 - are protein components of the SMN complex, integral to the formation of the core of snRNPs in the cytoplasm. The role of these proteins in the nucleus is less clear, however their increased interaction with SMNΔ7 suggests that SMNΔ7 is still incorporated into the SMN complex, most likely due to its ability to dimerise – albeit less effectively - with native, full-length SMN. (Le et al., 2005)

This methodology is ideal for lead generation and is particularly useful in the study of disease-relevant proteins when followed up with other modalities such as western blotting and immunostaining. Comparing the interactomes produced between a native protein and its disease-relevant mutant can allow for exploration of interactions gained and lost in pathology, as was performed for the SMN and SMNΔ7 proteins described herein. This data can then be leveraged via bioinformatics to further analyse pathways involved in the disease state, leading to the potential identification of clinically relevant future protein targets.

## Supporting information

Figure S1 and details of figure S2 and table S1

Figure S2 (interactive version of Fig 5)

Table S1: neuronal proteins

## References

Agha-Mohammadi, S., O’Malley, M., Etemad, A., Wang, Z., Xiao, X., & Lotze, M. T. (2004). Second-generation tetracycline-regulatable promoter: repositioned tet operator elements optimize transactivator synergy while shorter minimal promoter offers tight basal leakiness. J Gene Med, 6(7), 817–828. 10.1002/jgm.566

Branon, T. C., Bosch, J. A., Sanchez, A. D., Udeshi, N. D., Svinkina, T., Carr, S. A., Feldman, J. L., Perrimon, N., & Ting, A. Y. (2018). Efficient proximity labeling in living cells and organisms with TurboID. Nature Biotechnology, *36*(9), 880–887. 10.1038/nbt.4201

Cauchi, R. J. (2010). SMN and Gemins: ‘we are family’ … or are we?: insights into the partnership between Gemins and the spinal muscular atrophy disease protein SMN. BioEssays, 32(12), 1077–1089. 10.1002/bies.201000088

Chen, X., Wei, S., Ji, Y., Guo, X., & Yang, F. (2015). Quantitative proteomics using SILAC: Principles, applications, and developments. PROTEOMICS, 15(18), 3175–3192. 10.1002/pmic.201500108

Fedorova, S. A., & Dorogova, N. V. (2019). Protein trap: a new Swiss army knife for geneticists? [journal article]. Molecular Biology Reports. 10.1007/s11033-019-05181-z

Free, R. B., Hazelwood, L. A., & Sibley, D. R. (2009). Identifying Novel Protein-Protein Interactions Using Co-Immunoprecipitation and Mass Spectroscopy. Current Protocols in Neuroscience, *46*(1). 10.1002/0471142301.ns0528s46

Fuller, H. R., Mandefro, B., Shirran, S. L., Gross, A. R., Kaus, A. S., Botting, C. H., Morris, G. E., & Sareen, D. (2015). Spinal Muscular Atrophy Patient iPSC-Derived Motor Neurons Have Reduced Expression of Proteins Important in Neuronal Development. *Frontiers in cellular neuroscience*, 9, 506. 10.3389/fncel.2015.00506

Hopp, T. P., Prickett, K. S., Price, V. L., Libby, R. T., March, C. J., Pat Cerretti, D., Urdal, D. L., & Conlon, P. J. (1988). A Short Polypeptide Marker Sequence Useful for Recombinant Protein Identification and Purification. *Bio/Technology*, 6(10), 1204–1210. 10.1038/nbt1088-1204

Kimple, M. E., Brill, A. L., & Pasker, R. L. (2013). Overview of Affinity Tags for Protein Purification. Current protocols in protein science, 73(1). 10.1002/0471140864.ps0909s73

Le, T. T., Coovert, D. D., Monani, U. R., Morris, G. E., & Burghes, A. H. (2000). The survival motor neuron (SMN) protein: effect of exon loss and mutation on protein localization. Neurogenetics, 3(1), 7–16. 10.1007/s100480000090

Le, T. T., Pham, L. T., Butchbach, M. E., Zhang, H. L., Monani, U. R., Coovert, D. D., Gavrilina, T. O., Xing, L., Bassell, G. J., & Burghes, A. H. (2005). SMNDelta7, the major product of the centromeric survival motor neuron (SMN2) gene, extends survival in mice with spinal muscular atrophy and associates with full-length SMN. Hum Mol Genet, 14(6), 845–857. 10.1093/hmg/ddi078

Lefebvre, S., Bürglen, L., Reboullet, S., Clermont, O., Burlet, P., Viollet, L., Benichou, B., Cruaud, C., Millasseau, P., Zeviani, M., Le Paslier, D., Frézal, J., Cohen, D., Weissenbach, J., Munnich, A., & Melki, J. (1995). Identification and characterization of a spinal muscular atrophy-determining gene. Cell, 80(1), 155–165. 10.1016/0092-8674(95)90460-3

Lefebvre, S., Burlet, P., Liu, Q., Bertrandy, S., Clermont, O., Munnich, A., Dreyfuss, G., & Melki, J. (1997). Correlation between severity and SMN protein level in spinal muscular atrophy. Nature Genetics, *16*(3), 265–269. 10.1038/ng0797-265

Li, P., Meng, Y., Wang, L., & Di, L. J. (2019). BioID: A Proximity-Dependent Labeling Approach in Proteomics Study. Methods Mol Biol, 1871, 143–151. 10.1007/978-1-4939-8814-3_10

Liu, X., Abad, L., Chatterjee, L., Cristea, I. M., & Varjosalo, M. (2024). Mapping protein–protein interactions by mass spectrometry. Mass Spectrometry Reviews. 10.1002/mas.21887

Lo Sardo, F. (2023). Co-Immunoprecipitation (Co-Ip) in Mammalian Cells. Methods Mol Biol, 2655, 67–77. 10.1007/978-1-0716-3143-0_6

May, D. G., Scott, K. L., Campos, A. R., & Roux, K. J. (2020). Comparative Application of BioID and TurboID for Protein-Proximity Biotinylation. Cells, 9(5), 1070. 10.3390/cells9051070

Mezentsev, Y., Ershov, P., Yablokov, E., Kaluzhskiy, L., Kupriyanov, K., Gnedenko, O., & Ivanov, A. (2022). Protein Interactome Profiling of Stable Molecular Complexes in Biomaterial Lysate. International Journal of Molecular Sciences, 23(24), 15697. 10.3390/ijms232415697

Samavarchi-Tehrani, P., Samson, R., & Gingras, A.-C. (2020). Proximity dependent biotinylation: key enzymes and adaptation to proteomics approaches. Molecular & Cellular Proteomics, mcp.R120.001941. 10.1074/mcp.r120.001941

Shcherbo, D., Murphy, C. S., Ermakova, G. V., Solovieva, E. A., Chepurnykh, T. V., Shcheglov, A. S., Verkhusha, V. V., Pletnev, V. Z., Hazelwood, K. L., Roche, P. M., Lukyanov, S., Zaraisky, A. G., Davidson, M. W., & Chudakov, D. M. (2009). Far-red fluorescent tags for protein imaging in living tissues. Biochem J, 418(3), 567–574. 10.1042/bj20081949

Varderidou-Minasian, S., Verheijen, B. M., Harschnitz, O., Kling, S., Karst, H., van der Pol, W. L., Pasterkamp, R. J., & Altelaar, M. (2021). Spinal Muscular Atrophy Patient iPSC-Derived Motor Neurons Display Altered Proteomes at Early Stages of Differentiation. *ACS Omega*, 6(51), 35375-35388. 10.1021/acsomega.1c04688

Weimer, K., Zambo, B., & Gogl, G. (2023). Molecules interact. But how strong and how much? BioEssays, 45(6), 2300007. 10.1002/bies.202300007

