## Supplementary material for "Investigating the Interactomic Landscape of Survival Motor Neurons (SMN) and the SMN Δ7 truncated protein": Figure S1 and details of figure S2 and table S1

**Supplementary Information**

**S1: TurboID Lentiviral Plasmid Maps**

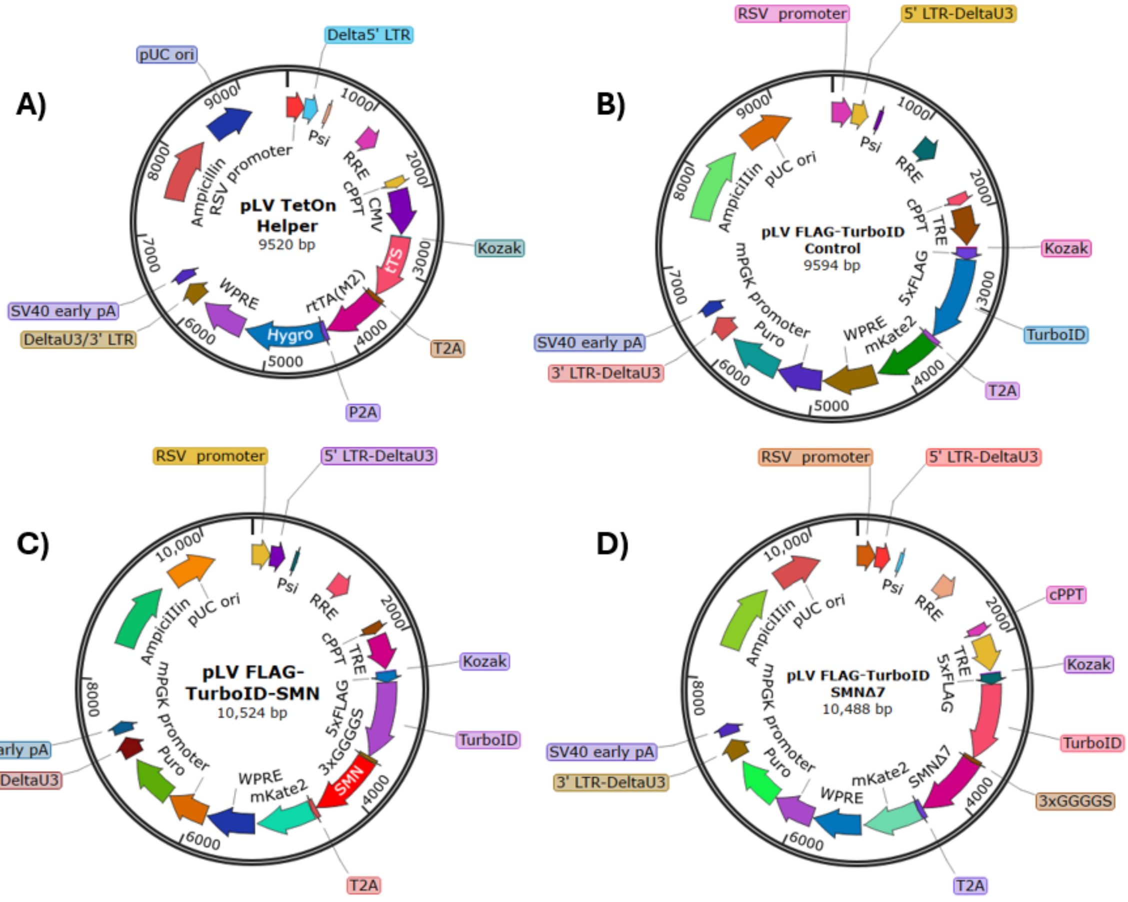


**Figure S1:** Map of the TetOn Helper plasmid. B) Map of LV Plasmid designed to express FLAG-tagged TurboID (control plasmid) C) Map of LV Plasmid designed to express FLAG-tagged TurboID-SMN. C) Map of LV Plasmid designed to express FLAG-tagged TurboID-SMNΔ7.

Note: These plasmids contain a TetOn system; The Tet Repressor protein, which cannot bind to the TetOn transcription elements when dox is absent, was coded into one plasmid (described above as the TetOn Helper plasmid) which must first be transduced into cells prior to TurboID plasmid transduction. This plasmid also encodes a silencer which binds to upstream promoter elements in the absence of doxycycline, preventing expression and reducing any leakage inherent in the system (Agha-Mohammadi et al., 2004).

**Figure S2** The Streptavidin Pulldown Data demonstrates Greater Interactome coverage than FLAG IP. Interactive versions of the plot in figure 5

**Table S1: Neuronally-enriched proteins (derived from the Human Protein Atlas, October 2024)**
